# 3D-surface reconstruction of cellular cryo-soft X-ray microscopy tomograms using semi-supervised deep learning

**DOI:** 10.1101/2022.05.16.492055

**Authors:** Michael C. A. Dyhr, Mohsen Sadeghi, Ralitsa Moynova, Carolin Knappe, Burcu Kepsutlu, Stephan Werner, Gerd Schneider, James McNally, Frank Noe, Helge Ewers

**Author notes:** equal contribution (Michael C. A. Dyhr, Mohsen Sadeghi).

## Abstract

Cryo-soft X-ray tomography (cryo-SXT) is a powerful method to investigate the ultrastructure of cells, offering resolution in the tens of nm range and strong contrast for membranous structures without requirement for labeling or chemical fixation. The short acquisition time and the relatively large volumes acquired allow for fast acquisition of large amounts of tomographic image data. Segmentation of these data into accessible features is a necessary step in gaining biologically relevant information from cryo-soft X-ray tomograms. However, manual image segmentation still requires several orders of magnitude more time than data acquisition. To address this challenge, we have here developed an end-to-end automated 3D-segmentation pipeline based on semi-supervised deep learning. Our approach is suitable for high-throughput analysis of large amounts of tomographic data, while being robust when faced with limited manual annotations and variations in the tomographic conditions. We validate our approach by extracting three-dimensional information on cellular ultrastructure and by quantifying nanoscopic morphological parameters of filopodia in mammalian cells.

## Introduction

All cells are filled with a dense and complex mixture of organelles and macromolecular structures that rely on nanoscale interactions to perform vital functions. To understand the organization and interaction of cellular organelles, the resolution of native cellular ultrastructure is essential. Electron microscopy (EM) is considered the gold standard for ultrastructural analysis. However, while much of our current understanding of cellular processes is based on EM methods, several limitations in sample preparation, volume coverage, acquisition speed and throughput remain. Cryo-soft X-ray microscopy (XRM) offers several advantages over EM approaches, but with the trade-off of somewhat lower resolution.

XRM takes advantage of the intrinsic different absorption contrast of elements. In the spectral region called the ‘water window’, carbon atoms absorb very strongly compared to oxygen atoms. This is of great relevance for biological specimens, where the natural contrast between carbon-dense material such as proteins and lipids and the aqueous environment can be exploited to visualize membranous organelles and small particles (e.g. nanoparticles^1^ or viruses^2^) at a resolution of 30 - 40 nm^3^. Moreover, specific organelles or structures can be distinguished based on their linear absorption coefficient, which is directly related to their chemical composition^4^. Unlike many EM methods, cryo-XRM samples require neither chemical fixation nor embedding or labeling. Instead, cells can be grown on gold TEM grids, plunge frozen and directly imaged in the transmission x-ray microscope. Furthermore, the penetration depth of soft x-rays is with 15 μm significantly deeper than what would be accessible by transmission electron microscopy techniques. This allows transmission microscopy and tomography of the entirety of the cellular volume in minutes without the need for physical sectioning^5,6^.

Cryo-soft X-ray tomography (SXT) yields projection images acquired at tilt angles within ±65°, which are then aligned and a tomographic reconstruction algorithm is then applied to produce a tomogram. In this paper, we use the term “tomogram” instead of “tomographic reconstruction” in order to avoid confusion with the reconstruction procedure performed by our network. This tomographic dataset composed of hundreds of image slices contains the 3D-ultrastructure of the cell that must then be extracted using segmentation to yield more quantifiable information, as for instance the size, organization and distribution of organelles in the cytosol^3^. In comparison to conventional EM methods, cryo-SXT covers a significantly larger volume in a shorter period of time and does not suffer from preparation artifacts such as section compression or shrinkage^7,8^. Fast-paced development of 3D-EM techniques towards larger volumes and imaging under cryogenic and non-cryogenic conditions has led to remarkable results in the ultrastructure of biological specimens^9–12^. Sample preparation and acquisition times of techniques such as focused ion beam (FIB) SEM are several orders of magnitude higher than those of SXT^13^, meaning that the throughput is generally low. In contrast, cryo-SXT can acquire a tomogram covering ∼1000 μm^3^ within less than 30 min and thus allows the comparison of changes in i.e. the organization of cellular organelles with statistically relevant amounts of data in 3D volumes as in the yeast cell cycle^14^, nanoparticle uptake^1^ and parasite infection^15^. Moreover, the compatibility of cryo-SXT with cryo-fluorescence microscopy or cryo-hard x-ray fluorescence microscopy allows the identification and visualization of specific cellular structures and processes^16,17^.

While volumetric data can be generated rather quickly in cryo-SXT, the data analysis is still time-consuming. Specifically, in the cryo-SXT workflow, manual segmentation remains the key bottleneck^18^. The complete manual segmentation of a dataset may take several days or even weeks, discouraging effort to segment the data beyond the structures of interest. As a result, subtle but potentially important changes in cellular ultrastructure may go unnoticed, even if that information was present in the original data. Automated or semi-automated segmentation using machine learning algorithms, especially artificial neural networks, hold the promise of overcoming this barrier^19–21^. Deep learning (DL) has already made great impact in microscopy, with promising applications in resolution enhancement^22–25^ and denoising^26^, missing data reconstruction^27^ and rapid phenotyping^28^. More specifically, first contributions have already been made for automated segmentation in various microscopy applications, including volume-EM^9,10,29,30^. Open source solutions offering basic functionality are also available^31–34^ and the design of dedicated novel networks will likely lead to further improvements. The design of methods suited for automated segmentation of cryo-SXT tomograms is an ongoing effort^35–39^ and faces inherent challenges such as limited depth of focus as well as the missing wedge problem^6^. The missing wedge refers to larger tilt angles that are missing in the 3D tomogram, for example in XRM, +/-65° instead of +/-90°. Reconstruction of data with a missing wedge introduces artifacts, such as blurring and apparent elongation in the Z-axis^40,41^. Addressing these challenges using DL image analysis requires a large pool of consistent, high-quality training data, which in turn requires extensive access to suitable cryo-soft X-ray microscopes for acquiring such data.

Deep convolutional networks remain the most popular approach for supervised learning of image segmentation tasks^42–44^, while other approaches such as learning weighted random walks have also been suggested^45^. Relying exclusively on supervised learning requires access to large amounts of manually annotated data. Moreover, in supervised training the network can overfit on the instrument(s) used in the training data and fail to generalize to a different imaging source. To address these challenges, we here introduce a deep convolutional network for semi-supervised learning of cryo-SXT image segmentation. We propose a new architecture that uses a combination of manually annotated as well as unannotated images for training. While the network learns to classify features of interest, such as membrane structures, in the tomographic image using the manual annotations, significantly larger amounts of unannotated data enable the network to learn a representation of various possible image features and imaging conditions, and thereby significantly reduce the amount of annotated images required to make reliable predictions. Once trained on both annotated and unannotated data, the network can be applied out-of-the-box for automatic segmentation and surface reconstruction of a variety of tomograms from different instruments without the need for parameter tweaking and retraining. The entire procedure requires minutes for whole-cell tomograms, offering a versatile tool to fully harness the structural information in cryo-soft X-ray tomographic datasets. We test our model on unseen data acquired at three different synchrotrons to demonstrate transferability and quantify a significant amount of subresolution 3D data to demonstrate the throughput of our technique.

## Results

### Semi-supervised deep neural network for cryo-SXT segmentation

We first needed to develop an end-to-end deep-learning network suited for the analysis of cryo-SXT datasets. After testing a large number of different network architectures, we determined that the significant variety of image features and the relatively low contrast in cryo-SXT images demand a very deep network with a large representation power.

Consequently, we constructed a convolutional feed forward network, (Figure 1, for details on network design and training, see *Methods*)). This network takes 2D slices of cryo-SXT tomograms, and processes them to two parallel outputs by using “image” and “annotation” decoders, respectively^19^. Besides width and height dimensions, an image processed by the network is assigned an extra “channel” dimension, which is one for the input grayscale images. The channel dimension contains parallel sets of information about an image, that are being processed simultaneously at each layer of the network. Since using residual learning allows the training of significantly deeper networks^46^, our proposed architecture heavily relies on residual blocks made up of convolutional layers with residuals concatenated in channel space, accumulating extended pixel-wise information.

**Figure 1:**
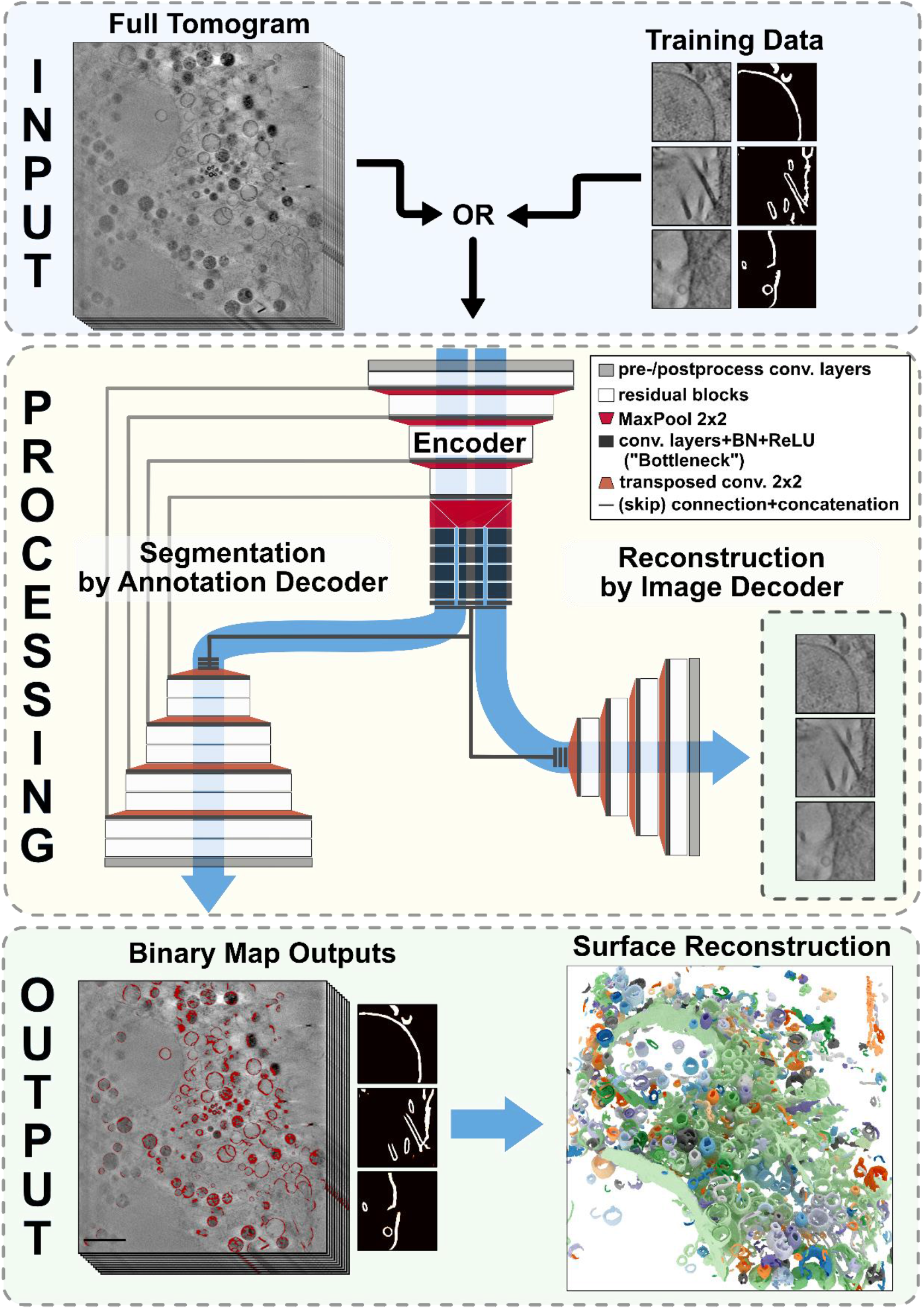
Processing workflow from raw tilt data to 3D volume rendering of cryo-SXT datasets. Top, INPUT: A full tomogram or manually annotated training data are used as input. Shown are a 0° tilt image of a representative cryo-soft X-ray tomographic dataset, showing a part of the cytosol of a plunge-frozen CRFK cell and examples of training data presented to the network. **Middle, Processing:** Schematic of the two-headed convolutional neural network. **Bottom, Output, from left:** Overlay of the tomographic slice shown at the top and the corresponding network output labels (in red). Three segmentation outputs for the training data input shown in the top right. 3D volume rendering of all network output labels from the entire tomographic reconstruction with the detected and reconstructed objects displayed in random colors. Scale bar: 2 μm.

Overall, our deep learning network contains 30 layers in the segmentation path (Fig. 1) with a total of ∼76 million trainable parameters. In order to facilitate semi-supervised training on cryo-soft X-ray tomograms, we have implemented a network architecture which allows for simultaneous learning of both image reconstruction and image segmentation. To improve the training procedure of the network, we have employed batch normalization in the majority of blocks, which leads to faster convergence by regularizing the mean and variance of brightness values across batches of image data^47^ (Fig. 1). The global bottleneck of the network introduces a cardinality of 3, i.e. three parallel convolutional paths with different kernel sizes (Fig. 1). This allows for higher representation power with less network depth, and prevents vanishing gradients during the training procedure^48^. The outputs of the bottleneck are concatenated and passed to the two decoders. The image decoder (Fig. 1) is tasked with reproducing the input image, while the annotation decoder performs the segmentation. The characteristic skip connections of the U-Net architecture^29,30^ are only introduced between the encoder and the annotation decoder (Fig. 1).

For the image segmentation branch, we have made hyperparameter choices similar to the original U-Net design in the number of down/up-sampling stages, kernel sizes, and number of channels. For other hyperparameters, such as the number of residual blocks in each stage, the cardinality of the bottleneck region, final layers in the decoders, and the learning rate, we have relied on extensive experimentation and manual selection of the combinations that resulted in best results, both qualitatively as judged by the final segmentation, and quantitatively as measured via pixel-wise accuracy and SSIM. After the training of the network is complete, full-sized slices can be processed in one-shot scans. In order to handle large image slices with limited memory, we have also implemented a procedure for partitioning images into overlapping chunks, and gathering the segmented labels subsequently.

The network was trained in a semi-supervised approach by two protocols: (i) we used a large number of slices from different tomograms to train the reconstruction path of the network without updating the weights in the segmentation decoder. This forces the encoder to learn a compressed representation transferable between different cell and tomography conditions. (ii) we used the smaller number of manually segmented inputs to train both paths simultaneously in order to reproduce the image and its accompanying label, respectively. The training loop entails alternating between the two protocols until satisfactory convergence is achieved (**Fig. 1**).

We have used a pixel-wise accuracy score that measures the percentage of pixels correctly predicted, respectively within 1% margin for segmented labels, and within one standard deviation of brightness for reconstructed images. After 700 training iterations, the image reconstruction path achieves a pixel-wise accuracy of > 80%. Reconstructing images also entails reproducing the background noise. Because the network lacks any stochastic input for noise sampling, the background noise is implicitly approximated (**Fig. 1** and **Fig. S1**). The segmentation path reproduces manual labels with pixel-wise test accuracy of > 96%.

### Evaluation of segmentation speed, quality and robustness

Using a GeForce RTX 3090 graphics card with 24 GB of graphics memory, and using the already trained network, automated segmentation took less than 5 minutes for a tomogram with 350 slices of 1324×1284 pixel images or 8 minutes for a tomogram of 500 similarly-sized slices, respectively. The performance was limited by the necessary slicing of large images into overlapping chunks suitable for GPU processing,, which is ultimately constrained by graphics card memory, and the resulting repeated data communications to and from the graphics card. Using more memory would thus further enhance the reported performance. A human expert user classified the automatically generated labels and filled in missing features into a representative set of 59 slices from 3 different tomograms. By comparing the pixels in each category, we found that out of all labeled pixels per slice, an average of 69.8% (s.d. = 10.4) were *true positives*, i.e. correctly labeled by the network and 27.7% (s.d. = 10.8) were *false negatives*, i.e. structures additionally labeled by the human user. 1.3 % (s.d. = 2.2) and 1.2 % (s.d. = 2.1) were *false positives* or labeled *reconstruction artifacts*, respectively (**Fig. 2**).

**Figure 2:**
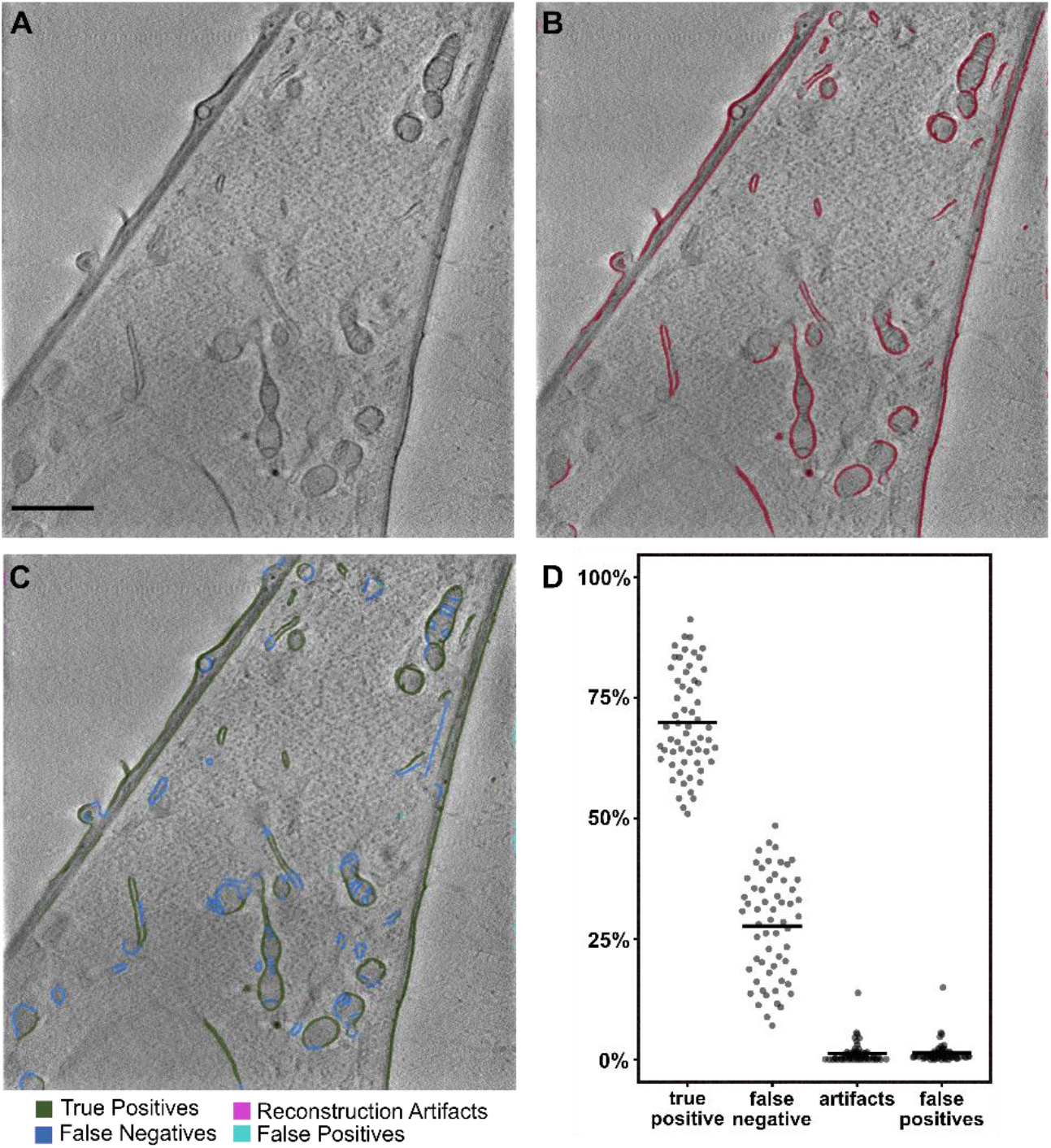
Quantification of Image Segmentation Robustness and Accuracy. **A)** A representative example D slice from the 3D tomographic image stack that was analyzed. **B)** Overlay of the example slice in A with the labels automatically produced by the deep network. **C)** The labels produced by the network were manually filled in on a series of randomly selected single slices and classified into true positives (green), reconstruction artifacts (pink), false positives (cyan). Manually added labels (false negatives) are shown in blue. **D)** The network performance was evaluated based on the percentage of surface pixels per class for each analyzed slice. Bars indicate the mean percentage. Scale bar: 2μm.

Since SXT requires synchrotron radiation, it can at this time only be performed at a few institutions worldwide. Thus, to be widely applicable, our method should generalize to data acquired form other synchrotrons. To test if this is the case, we applied our trained deep-learning network on X-ray microscopy datasets obtained from two other synchrotrons. We used a dataset of A549 lung cancer cells incubated with nanoparticles for endocytosis that was imaged on the MISTRAL beamline at the ALBA synchrotron (Barcelona, Spain; **Fig.S3**)^1^, and a dataset of mock-infected U2OS cells imaged on the B24 beamline at Diamond Light Source (Didcot, UK)^16^ (**Fig. S4**). The data provided by the Diamond Light Source were acquired with a different X-ray microscope objective with somewhat lower resolution but higher depth of field compared to the objectives used for all the other data (40 nm vs. 25 nm zone plates). Despite the different experimental conditions, both datasets were readily segmented by our trained network in seconds and identified a vast majority of membranous organelles inside the cytosol. These results demonstrate the utility of our deep network for data generated at cryo-soft x-ray microscopes from other synchrotron facilities and under different imaging and cell-biological conditions.

### Using deep learning to quantify filopodia morphology

Next, we asked whether our automated segmentation would allow for medium-throughput, high-resolution image analysis. As a ubiquitous example, we investigated cellular filopodia, which are abundant, sub-resolution plasma membrane protrusions, whose spatial dimensions are inaccessible by 2D-imaging. These elongated, submicron-thick, actin-filled structures play an important role in cellular migration and spreading of cells^49^. We obtained 3D images of 59 filipodia from three different cells, which we then quickly segmented, reconstructed and analysed (**Fig. 3A**).

**Figure 3:**
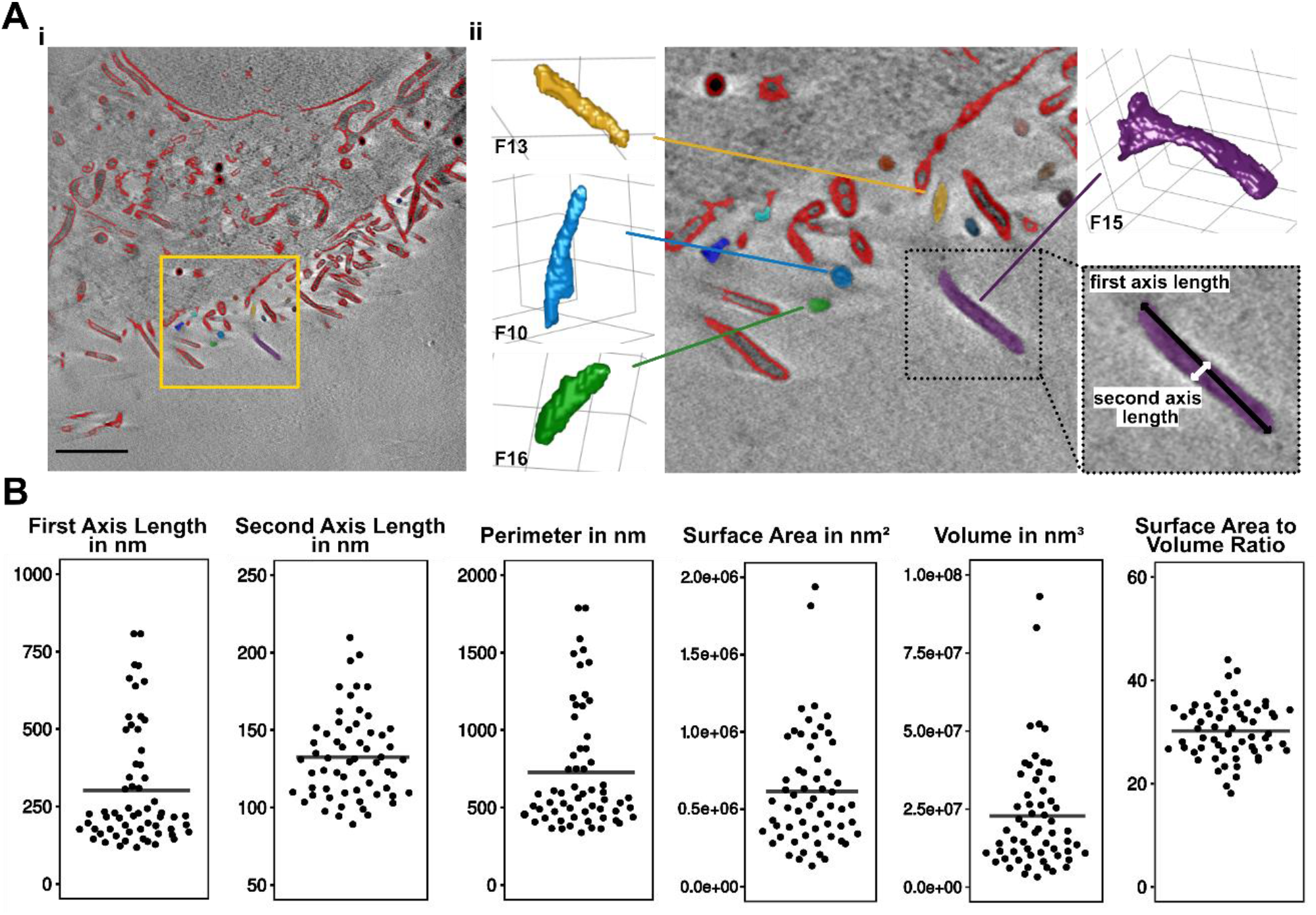
Quantification of filopodial 3D structure. **A)** Representative tomographic slice of an analysed dataset, overlaid with the automatically generated labels in red (i). The yellow box in *i* denotes the area shown up close in *ii*. A selection of individual filopodia used for analysis are shown in isosurface renderings, illustrating the general rod-like morphology of the filopodia. The box with dashed outline illustrates the orientation of first and second axis lengths on a representative filopodium. **B)** Dot-plots illustrating the mean first and second axis lengths and perimeter of the 2D cross-sections of the filopodia, as well as the 3D surface area, volume and surface area to volume ratio of all of the filopodia analysed in this study. The bar denotes the mean of the parameter across the population (n = 59). Scale bar: 2 μm.

As expected, we found that the spatial orientation of the filopodia affected their analysis in 2D-tomogram slices. Filopodia crossing a lateral tomographic slice at a low angle would be observed as a very elongated ellipse (i.e. having a long first axis length, **Fig. 3A**). Consistent with this observation, we noticed a high variation of the first axis length of the filopodia cross-section (302 ± 189 nm). The second axis length, more robustly reflecting filopodial diameter, was with 132 ± 28 nm generally more homogeneous. The diameter of filopodia is thus essentially constant but 3-dimensional orientation prevents accurate measurements of volume, surface area and perimeter from 2D cross-sections. It is thus essential to analyze entire filopodia from 3D-reconstructions.

Our fast deep learning-based reconstruction provided the basis for swift 3D analysis as well, in which we found a major axis length of 966 ± 394 nm by fitting an ellipsoid to the segmented data. The 3D-second axis length also had a higher error with 320 ± 141 nm (**Fig. 3B**). We found that the average volume and surface area of filopodia was highly variable, with 2.2 ·10^7^ ± 1.7 ·10^7^ nm³ and 6.1 ·10^5^ ± 3.6 ·10^5^ nm² respectively. However, the surface area to volume ratio was consistent with 30.2 ± 5.3 μm^-1^ (or 0.3021 ± 0.0534 nm^-1^, respectively) for the entire population, indicative of a consistent shape of all filopodia. Taken together, we concluded that the combination of SXT with DL image analysis allowed for fast reconstruction and analysis of small subresolution 3D structures.

## Discussion

In this work, we have developed a new deep neural network that allows for rapid and robust segmentation and surface reconstruction of three-dimensional cryo-SXT data. On average, it recognizes 70 % of the feature pixels in a cryo-soft X-ray tomogram within 10 minutes or less, regardless of the complexity of the dataset. Hence, even highly complex datasets such as mammalian cells can be segmented in a very short period of time. Even a semi-manual segmentation requires more than an order of magnitude more hands-on time by the user. We purposefully chose a more cautious automated segmentation approach with the deep network, since correcting false labels will be much more time-consuming than filling in missing labels. This suggests that future iterations with additional training data may recognize significantly more features.

Based on our segmentation evaluation, only about 1.8% or 1.6% of the automatic label pixels were either false positives or reconstruction artifacts on average per slice. These erroneous labels could at most times easily be distinguished from cellular features, since they were either localized in extracellular space and/or on the direct boundaries of the reconstructed volume. The appearance of these errors can be further reduced prior to classification without loss of information by cropping the dataset by a few pixels on the boundaries to avoid most reconstruction artifacts and by using a rough mask to differentiate the cellular volume from extracellular space to reduce the presence of false positives. Figure S2 illustrates examples of the falsely labeled reconstruction artifacts and false positives that were the most common in our datasets.

It is possible to perform the automatic segmentation with additional filtering to remove small objects, or to perform several iterations of erosion, leading to increasingly thinner contours and better object separation, as well as increasing the rate of false negatives. Depending on the intended use of the segmentation, i.e. rapid inspection vs. accurate analysis, as well as the size of the smallest objects of interest, tweaking of filtering and erosion settings may be useful for optimal results.

### Filopodia Analysis

To illustrate the utility of our deep network, we quantified several spatial parameters of a high number of filopodia. We found that 2D-analysis is not sufficient for the accurate determination of critical parameters, but 3D analysis after fast reconstruction through our algorithm allows for the swift determination of filopodial volume and surface area. These values have a high standard deviation, since the measured population represents filopodia in various states of extension and retraction of these highly dynamic and often transient structures^50^. The measurement of filopodial dimensions is due to their important role in cell migration and cancer of high interest ^49,51–54^. Our method will allow for a fast and convenient readout at the nanoscale and opens new avenues of research on filopodial growth.

### Advantages of our deep learning approach

Our network provides several key advantages for segmenting cryo-SXT data:

Using this network, a segmentation can be produced in less than ten minutes on a modern GPU. This dramatically simplifies and accelerates the data analysis step to the point where data can already be analyzed at the synchrotron during the acquisition of the data. This saves not only a significant amount of time, but also enables researchers to learn from their data faster and potentially adjust critical sample parameters before acquiring the next datasets, thereby increasing the quality of the data they obtain. This is especially important since cryo-SXT is predominantly available at synchrotrons, which only allows for limited amounts of measurement slots.

The manual input required for obtaining a 3D-segmentation of the data is significantly reduced, saving time and providing more objective segmentation results. The time required for a full segmentation of a dataset is considerable, so that users may focus their segmentation efforts on only a part of their data, looking for an expected result. An unbiased, automated segmentation can help in revealing more subtle or unexpected changes in the cells under the experimental conditions. At the same time, it becomes possible to generate statistically relevant amounts of data in a reasonable timeframe. This allows to take advantage of the full potential of cryo-SXT in indiscriminately visualizing all carbon-dense cellular structures, such as membranous and cytoskeletal structures.

Our network can be retrained with new training data. This should allow to improve the network’s performance under specific tomographic reconstruction conditions, or when different cell types other than adherent mammalian cells are being investigated. Thanks to the high processing speed and accuracy, it is a lot faster to generate a high number of training data based on preliminary segmentations, instead of manually segmenting a comparable amount of data. This can also be helpful for experimenting with other deep-learning tools.

The output data of the network can be exported as common file formats, including .TIFF format and thus easily imported into a variety of programs commonly used by biologists, such as ImageJ or Microscopy Image Browser (MIB). This ensures an easier integration into the user’s individual image processing and analysis workflow.

### Remaining Challenges

A few issues remain to be addressed in the future. While cryo-SXT as a method allows for rapid acquisition of a large portion of a cell, this imaging technique is still predominantly available at a few synchrotron facilities worldwide. Laboratory-based cryo-soft x-ray microscopes are under development but are currently not capable of achieving results of comparable quality. The limited accessibility of this technique, together with the diversity of applications and generated datasets results in a scarcity of comparable training data for effective training, which was one of the central challenges of this work.

The interior ultrastructure of mammalian cells is often highly complex, with many organelles in close proximity or even in direct contact with each other. This high degree of connectivity among the organelles poses a problem of object separation. This network was designed to rapidly and objectively segment all membranous structures of the cell. A user interested in separately quantifying one organelle from another must still invest manual effort in classifying and separating the objects (i.e. the organelles) from each other. The high speed and accuracy of our automated segmentation will now help to generate sufficient training data for such a classification network, which will learn to distinguish the major organelle types present in mammalian cells and assist in separating objects.

Finally, our automated procedure can be further improved to address the complications introduced by the *missing wedge problem*. This problem arises from incomplete rotation of a tomographic specimen, and cannot be avoided when imaging specimens grown on a flat substrate. Reconstruction of data with a missing wedge leads to blurring, elongation and ray artifacts which may obscure actual cellular features. As a consequence, cellular features distorted by missing-wedge artifacts become very difficult to discern, even for the human eye. Such cases formed only a small part of our training data space, and so our network is not yet sufficiently well trained to compensate for these artifacts. This will be overcome as the network is exposed to more and more data which suffer from such artifacts, and so with time the network will learn to more accurately segment features obscured by missing-wedge artifacts.

## Conclusion

In conclusion, we present here a convolutional neural network that dramatically reduces the manual effort and time required for segmentation of cryo-soft x-ray tomographic datasets. We show that the network recognizes about 70% of the features present in different datasets correctly and thus allows for an instant assessment of the dataset. This will allow soft X-ray microscope users to analyze their data more rapidly, efficiently and objectively, and thereby optimize their microscope time, which is constrained by the heavy demand for these instruments at synchrotron beamlines. Our deep network can segment the entire tomographic reconstruction within ten minutes or less without any manual user input. This allows users to assess their data as 3D-renderings during their experiment and as a result, they can make more informed choices about the use of their very limited beamtime and address their biological questions more efficiently. Furthermore, our work opens the door to the generation of statistically relevant amounts of data on large cytosolic volumes from many cells. This will be of great importance in the quantitative investigation of the organization and reorganization of the cellular organelle complement.

## Methods

### Sample preparation

Crandell-Rees Feline Kidney (CRFK) cells were cultivated in DMEM with 4,5 g/l D-glucose, supplemented with 2 mM glutamine and 10 % FCS inside a humidified incubator at 37 °C. For preparing specimens, Quantifoil R2/2 AuG200F1 finder or Au-HZB-2 grids (Quantifoil Micro Tools GmbH, Germany) were first plasma-cleaned, then sterilized in 70 % ethanol and placed inside empty wells of a 24-well plate. After letting the remaining ethanol evaporate, fresh full medium was added and subsequently CRFK cells were seeded, such that they had reached a confluency of 70 - 80 % the next day, when the cells were either fixed for two hours at room temperature in freshly prepared 2 % glutaraldehyde or directly plunge-frozen in liquid ethane. The frozen specimens were stored in liquid nitrogen until imaging.

### Cryo-soft x-ray tomography and tomographic reconstruction

The acquisitions were performed at the U41 beamline at the BESSY II synchrotron facility. Tilt images ranging from ± 60° (round Quantifoil grids) or ± 65° (HZB-2 grids) with an increment of 1° were acquired at 510 eV with a 25 nm Fresnel Zone Plate (FZP). Tracking, alignment and tomographic reconstruction were performed using the Simultaneous Iterative Reconstruction Technique (SIRT)^55^ in IMOD^56^, using gold nanoparticles as fiducial markers.

### Deep network for image segmentation

The architecture shown in Fig. 1 is an end-to-end convolutional feed forward network, taking 2D slices of cryo-SXT tomograms, and processing them to two concurrent outputs, respectively from “image” and “annotation” decoders^19^. Beside the familiar width and height dimensions, an image that is being processed by the network is assigned an extra “channel” dimension, holding an array of brightness information per pixel. Our proposed architecture heavily relies on residual blocks made up of convolutional layers with residuals concatenated in channel space, accumulating extended pixel-wise information. Using residual learning allows the training of significantly deeper networks^46^. Convolutional layers of the network apply a convolution with a learnable kernel plus an additive bias, followed by the nonlinear activation function *g*. Thus, for image *X* being transformed by the network, *X*_*n*+1_ = *g*(*W*_*n*_ * *X*_*n*_ + *b*_*n*_), with *W*_*n*_ and *b*_*n*_ being the trainable kernel and bias, respectively. We have used Rectified Linear Unit (ReLU) as the activation function *g* throughout the network. Each individual convolution step in the network is composed of mini “bottle-neck” convolutional blocks, as described by He *et al*.^46^. In these blocks, the number of image channels is reduced by a convolution step with a kernel of size 1×1, followed by the main convolution with a 3×3 kernel, and a mapping back to the desired output channels by another 1×1 kernel. This setup reduces the number of trainable parameters in the network, while enhancing the abstraction learned by each convolution step.

### Data pipeline

The manually annotated data consists of 79 pairs of raw 1324 × 1284 pixel slices from a single tomogram and their corresponding manually fully annotated binary labels. For semi-supervised training, these data are complemented with 513 unannotated slices from different tomograms, for which manual labels are not produced. These unannotated data are used for training the image-reconstruction branch of the network. The network is trained on 256 × 256 pixel images, or image/label pairs, depending on if the input data is taken from the set with manual labels or unannotated images. We designed an image augmentation procedure which randomly crops the input slices into these smaller images, while additionally applying random 90° rotations and horizontal/vertical reflections. We have implemented the entire data preparation and augmentation pipeline using TensorFlow Dataset API, which benefits from automated GPU processing, hence offering uninterrupted training of the network while augmented data is prepared on-the-fly. The stochastic manner in which the training data are prepared and fed to the network results in a more robust prediction as well as less chance of over-fitting. The original data are split 90 % - 10 % between training and hold-out test sets before the augmentation, with the quality of image reconstruction and the accuracy of label prediction verified on the test set.

### Training and optimization

For the decoder performing image segmentation, we have used the pointwise Huber loss between manual (*Y*) and predicted (*X*) annotations^57^,

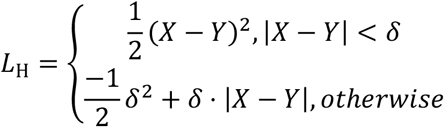

This function provides the robustness of l_1_-norm with constant gradients, with well-behaving near-zero gradients similar to the l_2_-norm. We found Huber loss to produce the sharpest segmentations with minimal artefacts. For the image reconstruction decoder, we have used as the loss function a combination of the Huber function and the Structural Similarity Index (SSIM), proposed by Wang and Sheikh^58^,

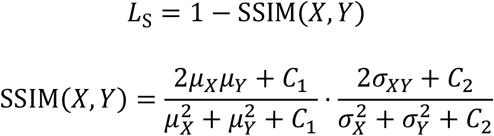

This choice has been backed by the systematic study of Zhao *et al*. on the loss functions for image reconstruction^59^.

Gradients of the loss function with respect to trainable parameters are calculated in the forward pass and the optimizer algorithm updates the parameters via backpropagation^25^. We monitor the training procedure by comparing the loss between training and test datasets. We stop the training loop when the test loss plateaus, which implies the network might begin to overfit to the training data.

In case the slice size results in memory shortage, we have also implemented a procedure for partitioning images into overlapping chunks, and gathering the segmented labels after the fact.

### 3D volumetric reconstruction

The two-dimensional outputs of the segmentation network provide a voxelized representation of the membranes in the tomogram. We have applied the well-established marching-cube algorithm to obtain surface reconstructions from the voxels^60^. The implementation from the SciPy package has been used for this purpose^61^. The PyVista python package has been used for processing of the 3D data^62^. Final renderings are achieved via the POV-Ray package^63^.

### Output export and import into software for downstream applications

The output labels generated by the deep network were exported as stacks of binary mask images in .TIF format. These stacks were then imported into *Microscopy Image Browser* ^64^. By thresholding, the binary labels could be quickly selected and used as a mask to be overlaid on the original input image. This result was saved as a. MODEL file and could be used for inspection of segmentation quality, label classification, manual complementation of labels and quantification of objects.

### Evaluating Segmentation Quality

In order to determine the quality of the automatic segmentation results, an expert user manually inspected the predicted labels of 59 randomly selected slices of 3 representative tomographic datasets, covering the entire range of the datasets by setting appropriate intervals between the slices. The labels generated by the network were overlaid with the corresponding, original slice and then classified into correct labels (a feature was being recognized correctly), false positives (a label was placed where no feature was present) and reconstruction artifacts. Cellular features that were missing in the network’s segmentation were manually labeled and classified as false negatives. The evaluation was performed by scoring the number of pixels of each category for each slice analyzed versus the total amount of labeled pixels. The inspection of slices, classification and evaluation was performed in Microscopy Image Browser^64^. Due to the missing wedge problem that arises in tomographic techniques with incomplete sample rotation and the limited depth of focus of the 25 nm FZP, some apparent features were present but too blurred for a human user to confidently trace the feature contour with a 4 pixel brush tool. These features were excluded from the analysis, since training a network to recognize such features would lead to a high increase of false positives.

### Filopodia Analysis

A total of sixty automatically labeled filopodia from three different datasets were selected for analysis. Filopodia had to be covered entirely by the reconstructed volume, i.e. filopodia protruding beyond the x, y, or z boundaries of the field of view were excluded, since 3D parameters cannot be conclusively determined from incomplete objects. Similarly, filopodia partially interrupted by e.g. reconstruction artifacts from gold fiducials were also excluded, since these artifacts prevent unambiguous labeling of underlying structures in tomographic datasets.

The automatically generated labels were inspected individually for each filopodium, tracing the entire structure from its base at the plasma membrane to the tip of the protrusion. While doing so, the object was manually separated from any adjacent labels, especially of the plasma membrane. Any open loops of the labels were closed and the contour’s shape filled, such that the volume and surface area values could be determined accurately. The isolated and filled objects were then quantified both as a series of 2D objects and as a fitted ellipsoid around the 3D object using the built-in quantification toolbox in *Microscopy Image Browser*^16^. For 2D quantification of each individual filopodium, the mean and standard deviation of all values in all slices of its component 2D objects were determined. The final quantification of all filopodia is based on the mean and standard deviation of all sixty individual means (2D) or 3D-parameters, respectively. For calculating the nanometer-scale values, an isotropic voxel size of 9.8 nm was used.

## Acknowledgements

This research has been funded by Deutsche Forschungsgemeinschaft (DFG) through grant SFB 958/Project A04, SFB 1114/Project C03, European Research Commission (CoG 772230 “ScaleCell”), the Berlin Institute for Foundations of Learning and Data (BIFOLD), through DFG project number 278001972 – TRR 186 and BMBF grant CLS9 COMPXRAY. Crandell-Rees Feline Kidney (CRFK) cells were a kind gift from Benedikt Kaufer lab. The acquisition of x-ray tomograms was performed at the U41-PGM1-XM beamline at BESSY II synchrotron facility unless otherwise indicated. The data collected for supplemental figure S3 was acquired at the MISTRAL beamline at ALBA synchrotron in collaboration with the beamline staff^1^. The data presented in supplemental figure S4 is based on the raw data of EMPIAR-10416 and EMPIAR-10417, which were kindly provided to us by the staff of B24 beamline at Diamond Light Source.

## Author Contributions

HE and FN acquired funding and conceptualized the study, MS and FN designed the deep neural network, MS developed the software and conducted network training, surface reconstructions, and image renderings. MD and BK performed experiments, image analysis and evaluations of the training efforts under guidance by JM, GS and HE. SW provided crucial technical guidance and support for cryo-SXT sample preparation and data collection. BK, CK and RM produced binary labels for the training data. MD, MS, JM, HE and FN wrote the paper.

---

**Table S1:**

|  | <u>Parameter</u> | <u>Mean ± SD (n=60)</u> |
| --- | --- | --- |
| 2D Analysis | Area Cross-section | 33155 ± 25244 nm<sup>2</sup> |
|  | First Axis Length | 302 ± 189 nm |
|  | Second Axis Length | 133 ± 28 nm |
|  | Perimeter | 727 ± 391 nm |
|  | Eccentricity | 0.80 ± 0.17 |
| 3D Analysis | Volume | 2.28 • 10<sup>7</sup> ± 1.77 • 10<sup>7</sup> nm<sup>3</sup> |
|  | Surface Area | 6.16 • 10<sup>5</sup> ± 3.66 • 10<sup>5</sup> nm<sup>2</sup> |
|  | Surface Area to Volume Ratio | 30.2 ± 5.3 μm<sup>-1</sup> |
|  | Major Axis Length | 966 ± 394 nm |
|  | Second Axis Length | 320 ± 141 nm |
|  | Third Axis Length | 206 ± 61 nm |

**Figure S1:**
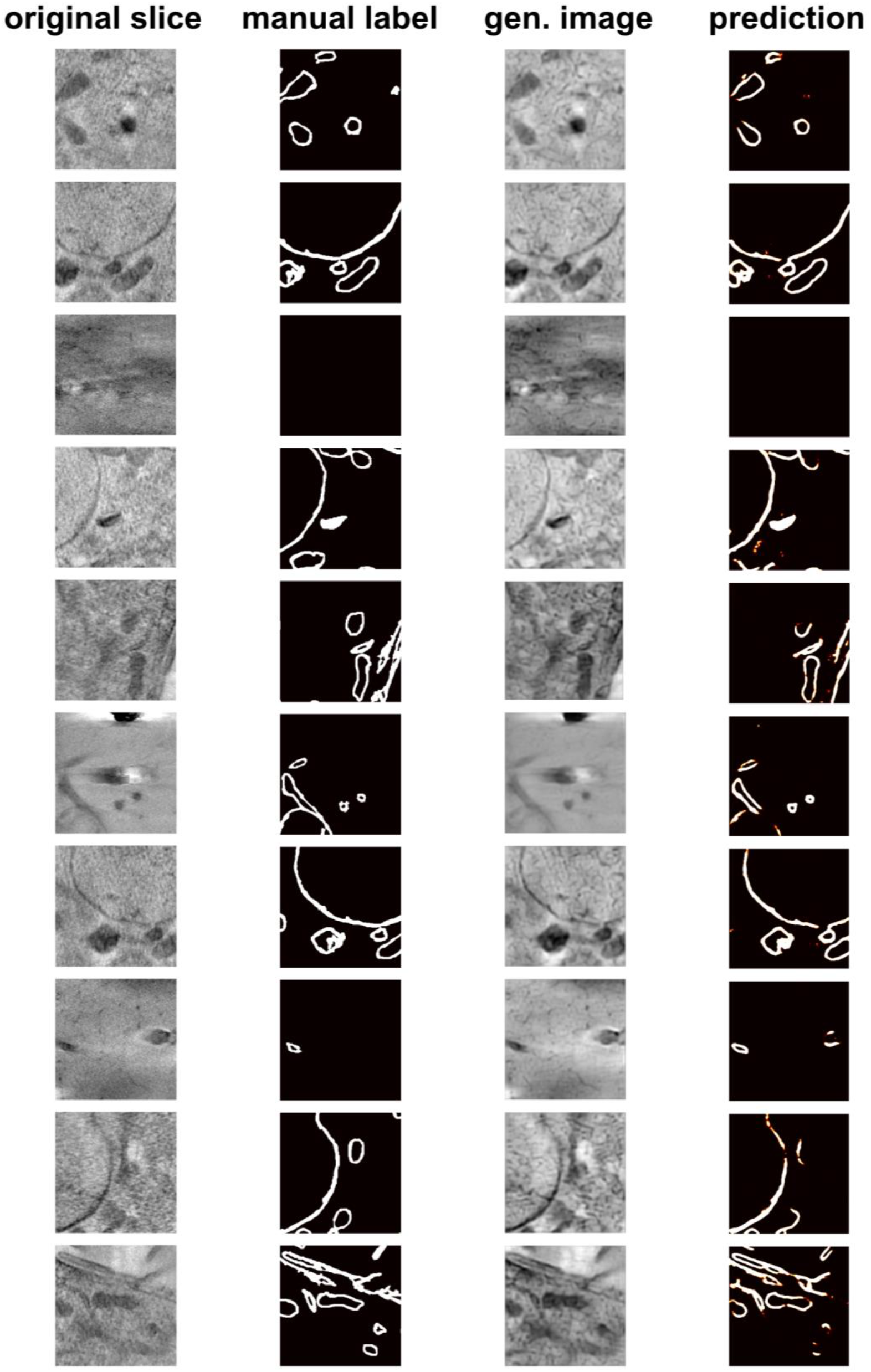
Training data. Shown are example images for input slices, their corresponding manual labels, as well as output images for the image decoder (generated images) and annotation decoder (prediction).

**Figure S2:**
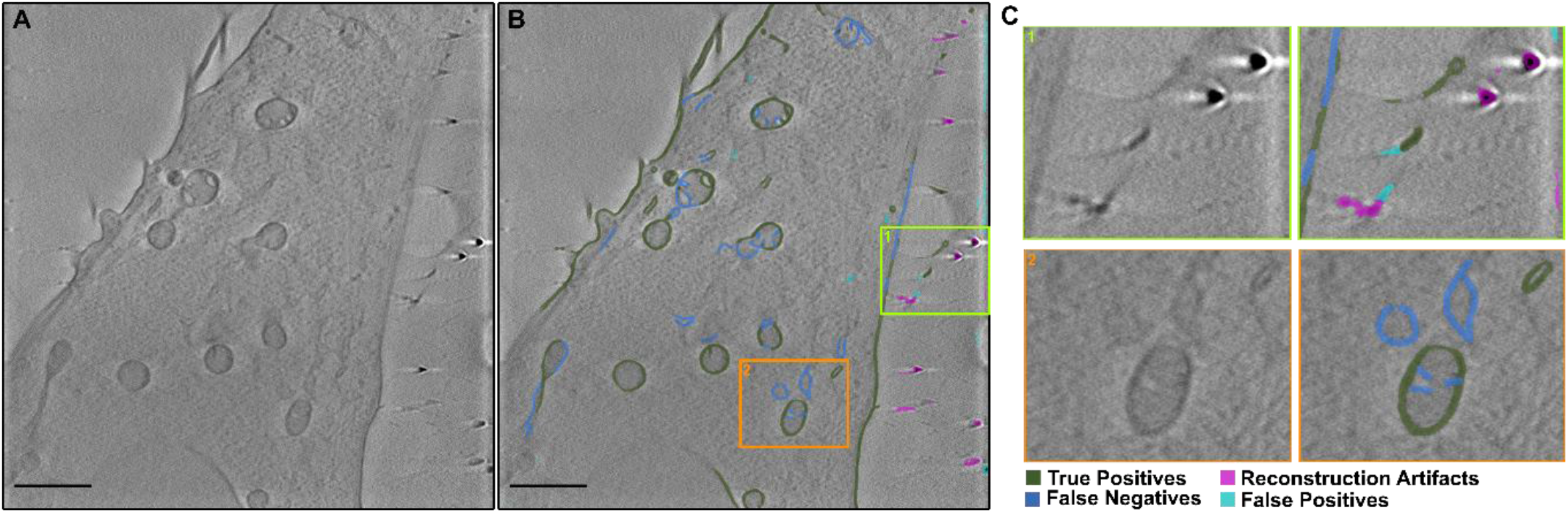
Examples for label categories. **A)** Raw input tomogram. **B)** Tomogram overlaid with labels categorized as “false positives”, “reconstruction artifacts”, “false negatives” and “true positives”. The green and orange boxes denote the close-ups in C. **C)** Close-ups of example regions, illustrating the label categories. Scale bars: 2μm.

**Figure S3:**
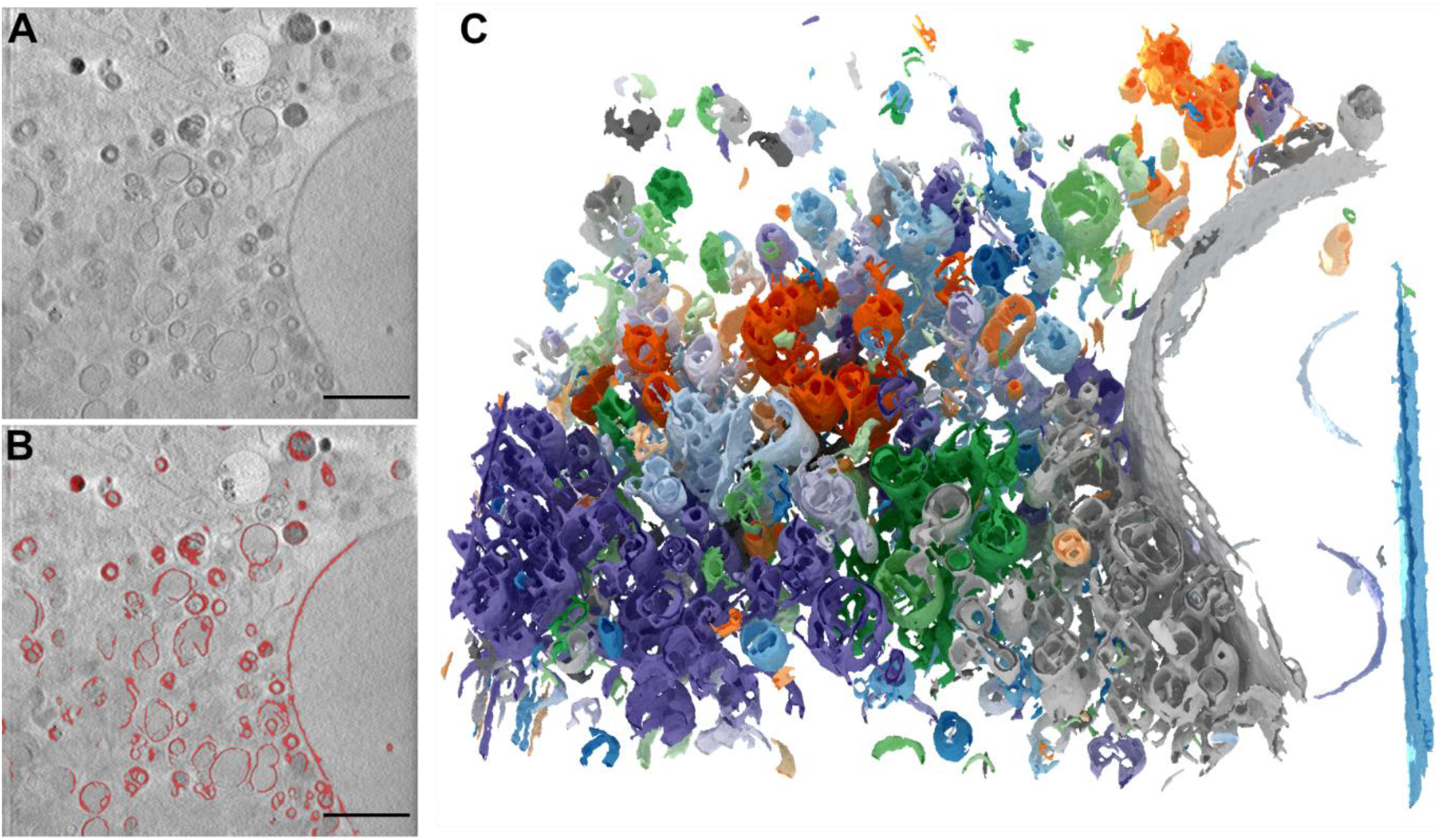
Reconstruction of Tomographic Data from the MISTRAL Beamline. **A)** Tomographic reconstruction of an A549 cell treated with PEI nanoparticles for 2 hours ^1^. **B)** Overlay of the raw tomogram (A) with automatically generated labels (red) by the deep network. **C)** 3D-rendering of the automated segmentation, objects shown in random colors. Scale bar: 2 μm.

**Figure S4:**
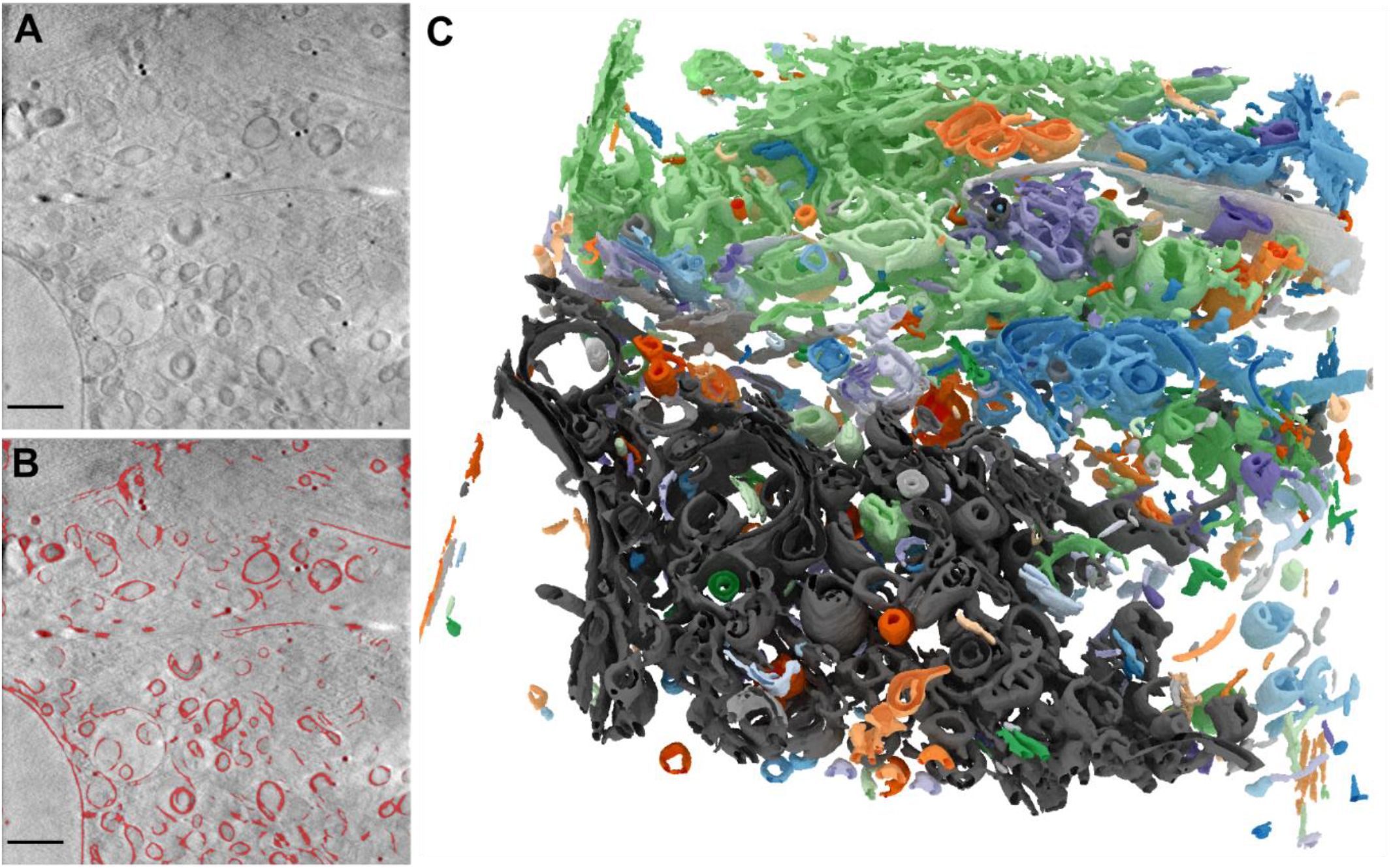
Reconstruction of Tomographic Data from the Diamond Light Source. **A)** Tomogram based on raw data for EMPIAR-10416 provided by Harkiolaki et al. (Diamond Light Source, UK). **B)** Overlay of the raw tomogram (A) with automatically generated labels (red) by the deep network. **C)** 3D-rendering of the automated segmentation, objects shown in random colors. Scale bars: 2μm.

## References

1. Kepsutlu, B. et al. Cells Undergo Major Changes in the Quantity of Cytoplasmic Organelles after Uptake of Gold Nanoparticles with Biologically Relevant Surface Coatings. ACS Nano 14, 2248–2264 (2020).

2. Chichón, F. J. et al. Cryo X-ray nano-tomography of vaccinia virus infected cells. J Struct Biol 177, 202–211 (2012).

3. Schneider, G. et al. Three-dimensional cellular ultrastructure resolved by X-ray microscopy. Nat Methods 7, 985–987 (2010).

4. Parkinson, D. Y., McDermott, G., Etkin, L. D., Le Gros, M. A. & Larabell, C. A. Quantitative 3-D imaging of eukaryotic cells using soft X-ray tomography. Journal of Structural Biology 162, 380–386 (2008).

5. Carzaniga, R., Domart, M.-C., Collinson, L. M. & Duke, E. Cryo-soft X-ray tomography: a journey into the world of the native-state cell. Protoplasma 251, 449–458 (2014).

6. Groen, J., Conesa, J. J., Valcárcel, R. & Pereiro, E. The cellular landscape by cryo soft X-ray tomography. Biophys Rev 11, 611–619 (2019).

7. Korogod, N., Petersen, C. C. & Knott, G. W. Ultrastructural analysis of adult mouse neocortex comparing aldehyde perfusion with cryo fixation. eLife 4, e05793 (2015).

8. Perktold, A., Zechmann, B., Daum, G. & Zellnig, G. Organelle association visualized by three dimensional ultrastructural imaging of the yeast cell. FEMS Yeast Research 7, 629–638 (2007).

9. Heinrich, L. et al. Whole-cell organelle segmentation in volume electron microscopy. Nature 1–6 (2021) doi:10.1038/s41586-021-03977-3.

10. Chen, M. et al. Convolutional neural networks for automated annotation of cellular cryo-electron tomograms. Nat Methods (2017) doi:10.1038/nmeth.4405.

11. Cortese, M. et al. Integrative Imaging Reveals SARS-CoV-2-Induced Reshaping of Subcellular Morphologies. Cell Host & Microbe 28, 853–866.e5 (2020).

12. Schaffer, M. et al. A cryo-FIB lift-out technique enables molecular-resolution cryo-ET within native Caenorhabditis elegans tissue. Nat Methods 16, 757–762 (2019).

13. Baena, V. et al. FIB-SEM as a Volume Electron Microscopy Approach to Study Cellular Architectures in SARS-CoV-2 and Other Viral Infections: A Practical Primer for a Virologist. Viruses 13, 611 (2021).

14. Uchida, M. et al. Quantitative Analysis of Yeast Internal Architecture using Soft X-ray Tomography. Yeast 28, 227–236 (2011).

15. Kapishnikov, S. et al. Unraveling heme detoxification in the malaria parasite by in situ correlative X-ray fluorescence microscopy and soft X-ray tomography. Sci Rep 7, 7610 (2017).

16. Kounatidis, I. et al. 3D Correlative Cryo-Structured Illumination Fluorescence and Soft X-ray Microscopy Elucidates Reovirus Intracellular Release Pathway. Cell 182, 515–530.e17 (2020).

17. Conesa, J. J. et al. Unambiguous Intracellular Localization and Quantification of a Potent Iridium Anticancer Compound by Correlative 3D Cryo X-Ray Imaging. Angewandte Chemie International Edition 59, 1270–1278 (2020).

18. Loconte, V. et al. Using soft X-ray tomography for rapid whole-cell quantitative imaging of SARS-CoV-2-infected cells. Cell Reports Methods 1, 100117 (2021).

19. Goodfellow, I., Bengio, Y. & Courville, A. Deep Learning. (MIT Press, 2016).

20. Carleo, G. et al. Machine learning and the physical sciences. Rev. Mod. Phys. 91, 045002 (2019).

21. Noé, F., Tkatchenko, A., Müller, K.-R. & Clementi, C. Machine Learning for Molecular Simulation. Annu. Rev. Phys. Chem. 71, 361–390 (2020).

22. Rivenson, Y. et al. Deep learning microscopy. Optica, OPTICA 4, 1437–1443 (2017).

23. Nehme, E., Weiss, L. E., Michaeli, T. & Shechtman, Y. Deep-STORM: super-resolution single-molecule microscopy by deep learning. Optica, OPTICA 5, 458–464 (2018).

24. Wang, H. et al. Deep learning enables cross-modality super-resolution in fluorescence microscopy. Nat Methods 16, 103–110 (2019).

25. Wang, Z., Chen, J. & Hoi, S. C. H. Deep Learning for Image Super-Resolution: A Survey. IEEE Transactions on Pattern Analysis and Machine Intelligence 43, 3365–3387 (2021).

26. Fang, L. et al. Deep learning-based point-scanning super-resolution imaging. Nat Methods 18, 406–416 (2021).

27. Chai, X. et al. Deep learning for irregularly and regularly missing data reconstruction. Sci Rep 10, 3302 (2020).

28. Sommer, C., Hoefler, R., Samwer, M. & Gerlich, D. W. A deep learning and novelty detection framework for rapid phenotyping in high-content screening. MBoC 28, 3428–3436 (2017).

29. Ronneberger, O., Fischer, P. & Brox, T. U-Net: Convolutional Networks for Biomedical Image Segmentation. in Medical Image Computing and Computer-Assisted Intervention –MICCAI 2015 (eds. Navab, N., Hornegger, J., Wells, W. M. & Frangi, A. F.) 234–241 (Springer International Publishing, 2015). doi:10.1007/978-3-319-24574-4_28.

30. Falk, T. et al. U-Net: deep learning for cell counting, detection, and morphometry. Nat Methods 16, 67–70 (2019).

31. Gómez-de-Mariscal, E. et al. DeepImageJ: A user-friendly environment to run deep learning models in ImageJ. Nat Methods 18, 1192–1195 (2021).

32. von Chamier, L. et al. Democratising deep learning for microscopy with ZeroCostDL4Mic. Nat Commun 12, 2276 (2021).

33. Belevich, I. & Jokitalo, E. DeepMIB: User-friendly and open-source software for training of deep learning network for biological image segmentation. PLoS Comput Biol 17, e1008374 (2021).

34. Berg, S. et al. ilastik: interactive machine learning for (bio)image analysis. Nat Methods 16, 1226–1232 (2019).

35. Plautz, T. et al. Progress Toward Automatic Segmentation of Soft X-ray Tomograms Using Convolutional Neural Networks. Microscopy and Microanalysis 23, 984–985 (2017).

36. Ekman, A., Chen, J.-H., Dermott, G. M., Gros, M. A. L. & Larabell, C. Task Based Semantic Segmentation of Soft X-ray CT Images Using 3D Convolutional Neural Networks. Microscopy and Microanalysis 26, 3152–3154 (2020).

37. Luengo, I. et al. SuRVoS: Super-Region Volume Segmentation workbench. Journal of Structural Biology 198, 43–53 (2017).

38. Nahas, K. L., Fernandes, J. F., Crump, C., Graham, S. & Harkiolaki, M. Contour, a semi-automated segmentation and quantitation tool for cryo-soft-X-ray tomography. 2021.12.03.470962 (2021) doi:10.1101/2021.12.03.470962.

39. Pennington, A. et al. SuRVoS 2: Accelerating Annotation and Segmentation for Large Volumetric Bioimage Workflows Across Modalities and Scales. Frontiers in Cell and Developmental Biology 10, (2022).

40. Koster, A. J. et al. Perspectives of Molecular and Cellular Electron Tomography. Journal of Structural Biology 120, 276–308 (1997).

41. Ercius, P., Alaidi, O., Rames, M. J. & Ren, G. Electron Tomography: A Three-Dimensional Analytic Tool for Hard and Soft Materials Research. Advanced Materials 27, 5638–5663 (2015).

42. Çiçek, Ö., Abdulkadir, A., Lienkamp, S. S., Brox, T. & Ronneberger, O. 3D U-Net: Learning Dense Volumetric Segmentation from Sparse Annotation. 1606.06650 [cs] (2016).

43. Lösel, P. D. et al. Introducing Biomedisa as an open-source online platform for biomedical image segmentation. Nat Commun 11, 5577 (2020).

44. Pelt, D. M. & Sethian, J. A. A mixed-scale dense convolutional neural network for image analysis. Proceedings of the National Academy of Sciences 115, 254–259 (2018).

45. Lösel, P. & Heuveline, V. Enhancing a diffusion algorithm for 4D image segmentation using local information. in Medical Imaging 2016: Image Processing vol. 9784 707–717 (SPIE, 2016).

46. He, K., Zhang, X., Ren, S. & Sun, J. Deep Residual Learning for Image Recognition. in 2016 IEEE Conference on Computer Vision and Pattern Recognition (CVPR) 770–778 (2016). doi:10.1109/CVPR.2016.90.

47. Ioffe, S. & Szegedy, C. Batch Normalization: Accelerating Deep Network Training by Reducing Internal Covariate Shift. 1502.03167 [cs] (2015).

48. Xie, S., Girshick, R., Dollár, P., Tu, Z. & He, K. Aggregated Residual Transformations for Deep Neural Networks. in 2017 IEEE Conference on Computer Vision and Pattern Recognition (CVPR) 5987–5995 (2017). doi:10.1109/CVPR.2017.634.

49. Jacquemet, G., Hamidi, H. & Ivaska, J. Filopodia in cell adhesion, 3D migration and cancer cell invasion. Current Opinion in Cell Biology 36, 23–31 (2015).

50. Bornschlögl, T. How filopodia pull: What we know about the mechanics and dynamics of filopodia. Cytoskeleton 70, 590–603 (2013).

51. Zhang, Q. et al. PTENε suppresses tumor metastasis through regulation of filopodia formation. EMBO J 40, e105806 (2021).

52. Gupton, S. L. & Gertler, F. B. Filopodia: The Fingers That Do the Walking. Science’s STKE 2007, re5–re5 (2007).

53. Gallop, J. L. Filopodia and their links with membrane traffic and cell adhesion. Semin Cell Dev Biol 102, 81–89 (2020).

54. Jacquemet, G. et al. FiloQuant reveals increased filopodia density during breast cancer progression. Journal of Cell Biology 216, 3387–3403 (2017).

55. Gilbert, P. Iterative methods for the three-dimensional reconstruction of an object from projections. Journal of Theoretical Biology 36, 105–117 (1972).

56. Kremer, J. R., Mastronarde, D. N. & McIntosh, J. R. Computer visualization of three-dimensional image data using IMOD. J Struct Biol 116, 71–76 (1996).

57. Huber, P. J. Robust Estimation of a Location Parameter. in Breakthroughs in Statistics: Methodology and Distribution (eds. Kotz, S. & Johnson, N. L.) 492–518 (Springer, 1992). doi:10.1007/978-1-4612-4380-9_35.

58. Wang, Z., Bovik, A. C., Sheikh, H. R. & Simoncelli, E. P. Image quality assessment: from error visibility to structural similarity. IEEE Transactions on Image Processing 13, 600–612 (2004).

59. Zhao, H., Gallo, O., Frosio, I. & Kautz, J. Loss Functions for Neural Networks for Image Processing. 1511.08861 [cs] (2018).

60. Lorensen, W. E. & Cline, H. E. Marching cubes: A high resolution 3D surface construction algorithm. in Proceedings of the 14th annual conference on Computer graphics and interactive techniques 163–169 (Association for Computing Machinery, 1987). doi:10.1145/37401.37422.

61. Virtanen, P. et al. SciPy 1.0: fundamental algorithms for scientific computing in Python. Nat Methods 17, 261–272 (2020).

62. Sullivan, C. B. & Kaszynski, A. A. PyVista: 3D plotting and mesh analysis through a streamlined interface for the Visualization Toolkit (VTK). Journal of Open Source Software 4, 1450 (2019).

63. POV-Ray - The Persistence of Vision Raytracer. https://www.povray.org/.

64. Belevich, I., Joensuu, M., Kumar, D., Vihinen, H. & Jokitalo, E. Microscopy Image Browser: A Platform for Segmentation and Analysis of Multidimensional Datasets. PLOS Biology 14, e1002340 (2016).

